# Molecular origins of asymmetric proton conduction in the influenza M2 channel

**DOI:** 10.1101/2022.07.31.502210

**Authors:** Themis Lazaridis

## Abstract

The M2 proton channel of influenza A is embedded into the viral envelope and allows acidification of the virion when the external pH is lowered. In contrast, no outward proton conductance is observed when the internal pH is lowered, although outward current is observed at positive voltage. Residues Trp41 and Asp44 are known to play a role in preventing pH-driven outward conductance but the mechanism for this is unclear. We investigate this issue using classical molecular dynamics simulations with stochastic proton hops. When all key His37 residues are neutral, inward proton movement is much more facile than outward movement if the His are allowed to shuttle the proton. The preference for inward movement increases further as the charge on the His37 increases. Analysis of the trajectories reveals three factors accounting for this asymmetry. First, the Asp44 trap the hydronium by strong electrostatic interactions. Secondly, Asp44 and Trp41 orient the hydronium with the protons pointing inward, hampering outward Grotthus hopping. The Trp41 add to the barrier by weakly H-bonding to potential H^+^ acceptors. Finally, for charged His, the H_3_O^+^ in the inner vestibule tends to get trapped at lipid-lined fenestrations of the cone-shaped channel. Simulations qualitatively reproduce the experimentally observed higher outward conductance of mutants. The ability of positive voltage, unlike proton gradient, to induce outward current appears to arise from its ability to bias H_3_O^+^ and the waters around it toward more H-outward orientations.

**Significance:** The M2 proton channel of influenza A, the best-studied viral ion channel and a proven drug target, conducts protons asymmetrically in response to a pH gradient. That is, protons flow inward when the external pH is low, but not outward when the internal pH is low. Experiments identified residues that play a role in this behavior, but how they do it has not been clear. This work identifies three molecular mechanisms that explain qualitatively the experimentally observed preference for inward conduction. These insights could improve our understanding of proton channels and possibly other key biological systems that exhibit vectorial proton transport.

## INTRODUCTION

Proton transport through membranes has a central role in bioenergetics (Nicholls and Ferguson 2002). All living cells store energy in the form of an electrochemical proton potential, which is subsequently converted to ATP. The proton potential is generated by integral membrane proteins that use light or chemical energy to move protons against their gradient. Although the structures of many proteins involved have been known for quite some time, the mechanisms of these processes are still not fully understood. Proton transport is also key to pH regulation mediated by proton channels (DeCoursey 2013). Therefore, determining the detailed mechanisms of proton transport would allow the understanding of biochemical systems that are fundamental to Life.

The M2 integral membrane protein of influenza A, the best studied member of the viroporin family (Nieva, Madan, and Carrasco 2012), acts as a proton channel that is activated by low external pH and allows the acidification of the virion in endosomes and the uncoating of the viral genome (Pinto and Lamb 2006). It also has a role in protecting acid-sensitive hemagglutinin during its transport to the plasma membrane (Grambas, Bennett, and Hay 1992). It is a single-pass membrane protein with a helical transmembrane domain that forms tetramers (Hong and DeGrado 2012). A key residue in the transmembrane domain is His37, which appears responsible for low pH activation and proton selectivity (Pinto, Holsinger, and Lamb 1992; Wang, Lamb, and Pinto 1995). Also conserved is a ring of Trp41 residues, one helical turn inward from the His37. This channel has attracted considerable experimental and theoretical work, with multiple structures available, from solid-state NMR (Sharma et al. 2010; Cady et al. 2010), solution NMR (Schnell and Chou 2008), and crystallography (Acharya et al. 2010; Thomaston et al. 2015, 2018).

Despite the extensive literature, certain aspects of M2 function are not completely understood. One of the most intriguing observations in M2 electrophysiology is the inability of the channel to conduct protons outward when the inside pH is lowered. Key to this behavior are residues Trp41 and Asp44, as their mutations led to observation of pH-driven outward currents (Tang et al. 2002; Ma et al. 2013). The D44N mutation is naturally observed in the Rostock strain and confers increased activity, which compensates for the lower acid stability of that strain’s hemagglutinin (Betakova, Ciampor, and Hay 2005). It has been proposed that the bulky Trp41 residues prevent access of the intraviral protons to the His ring and subsequent outward transport. However, in even the most “closed” M2 structures the Trp residues seem to leave enough space for H_3_O^+^ passage. In addition, it is unclear why a conformational change opening the gate would occur only when protons are flowing inward. Other attempts at explaining asymmetric conduction include the directionality of water wires (Thomaston et al. 2015) and the relative magnitude of free energy barriers on the two sides of the His ring (Liang et al. 2016). In recent work the loss of asymmetry in the D44N mutant was attributed to the more open structure at the C-terminus and the increased hydration below the His ring which allows activation from interior protons (Watkins, DeGrado, and Voth 2022). Finally, the question why voltage can drive outward current while ΔpH cannot has not yet been addressed.

Here we study the problem of conduction asymmetry in the M2 channel using a recently proposed algorithm for classical molecular dynamics (MD) simulations that allows “Grotthus” hopping of protons from one titratable site to another (Lazaridis and Hummer 2017). Using voltage as a biasing force to accelerate H^+^ translocation, we perform a large number of short simulations with a proton on either side of the Trp ring and measure the probability of crossing to the other side. We find asymmetry in proton crossing probability and identify the physical factors that determine this behavior in wild type and mutant channels.

## RESULTS

### Inward and outward crossing probabilities in the wild type channel

After equilibration of the 6BKK crystal structure in a POPC membrane (Fig. 1), a water molecule in the outer vestibule (just below Val27) or below the Asp44 was replaced by hydronium and positive or negative voltage was applied to drive it to the other side. The voltage magnitude was gradually increased until crossing events started to be observed. The protonation state of the His layer was set to either 0 or +2 (Q0, Q2). Five (sometimes ten) 100-ps simulations were performed in each case, with different initial velocities.

**Fig. 1.**
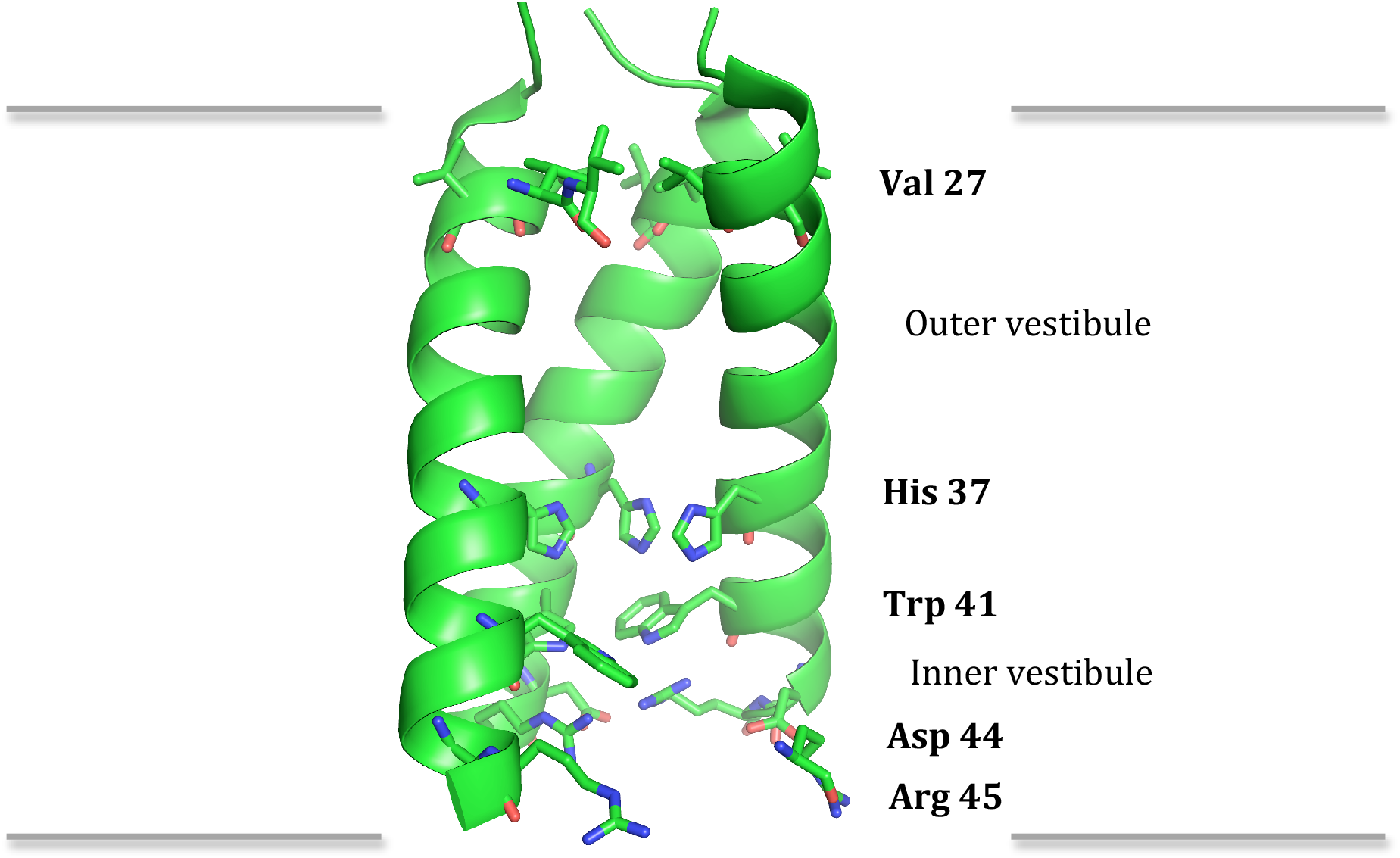
The structure of the M2 proton channel (pdb 6BKK) as embedded in a lipid bilayer. The front helix is omitted for clarity.

Two options were chosen for the inward simulations: allowing or not allowing the His to shuttle the proton by protonating and deprotonating. His protonation was not considered in the outward trajectories because they lie beyond the main barrier, which is the Trp41. Table 1 lists the percentage of trajectories that led to successful crossing of H^+^ to the other side of the Trp layer.

**Table 1.** Probability of crossing the Trp ring in the M2 channel as percentage of 5 or 10 100-ps trajectories where crossing is observed.

|  | Inward (no His titr) | Inward (His titr) | Outward |
| --- | --- | --- | --- |
| Q0-5V | 0% | 80% | 0% |
| Q0-8V | 40% | 80% | 30% |
| Q2-8V | 100% | 100% | 0% |

In the absence of His titration, neither inward nor outward conductance was observed in 100 ps at 5V. In the inward trajectories the H^+^ remains above the His layer, unable to find a way through other water molecules (the neutral His layer is quite dry). Allowing the His to protonate and deprotonate facilitates the movement of the proton. In some cases the H^+^ remains in one of the His or exchanges between the His for the duration of the simulation. In most cases it detaches and moves to the inner vestibule and beyond (note, however, that the kinetics of protonation and deprotonation of His is artificially accelerated in these simulations, see Methods). At higher voltage, conductance is observed in both directions even without His titration.

In the outward trajectories H_3_O^+^ is typically bound to the Asp44 side chains, sometimes escaping from one to bind to another (movie S1). Even though the Asp44 are salt-bridged to Arg45, they still have the ability to engage H_3_O^+^ with their free CO. In reality, close approach of a H^+^ to a deprotonated Asp could lead to Asp protonation, which is not allowed in the simulations (but see Discussion). However, the proximity of Arg45 probably keeps the Asp pK_a_ low, so we should expect at best transient protonation. Often the H_3_O^+^ lies between one Asp and 1 or 2 Trp residues and is oriented with the H inward (Fig. 2). In the trajectories with successful crossing at higher voltage the H_3_O^+^ is able to escape from the Asp and move outward through the Trp side chains. These results suggest a role for Asp44 as a proton trap.

**Fig. 2.**
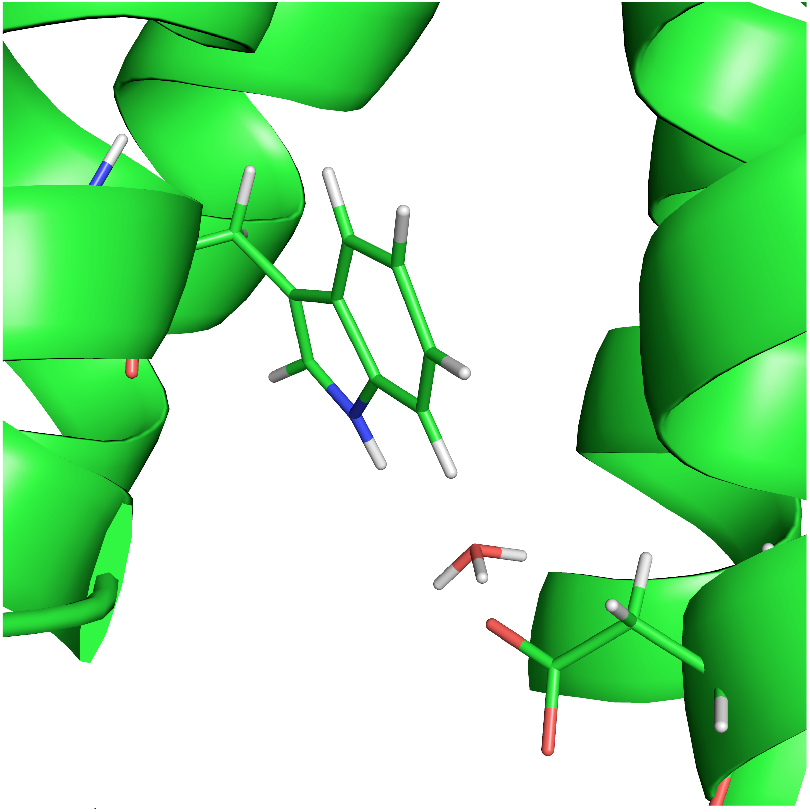
H_3_O^+^ trapped between Asp and Trp in M2 during an outward simulation at 5V.

The Q2-8V trajectories show an even more pronounced asymmetry, with or without titratable His. Inward conductance is facile with rapid inward movement of H_3_O^+^, which stays close to the center of the channel, oriented vertically (dipole parallel to the membrane), approaches one of the deprotonated His, and crosses easily the His and the Trp ring by Grotthus hopping through the His (if allowed) or other water molecules (movie S2). In the outward trajectories, H_3_O^+^ still interacts with the Asp but the repulsion from the His pushes it to the periphery of the channel from where outward movement is impossible (movie S3). It is remarkable how the same His charge repulsion has such disparate impact in the two directions.

In the Q2 trajectories the Trp gate is more open than in the Q0 trajectories. To quantify that, we measured the four indole Nε-Nε (“NN”) distances between adjacent Trp sidechains. In “closed” crystal structures, like the one used here, the Trp side chains are near each other with NN of 4.3-5.4 Å. This configuration is maintained in simulations of the neutral His state, but in the Q2 simulations the NN distances increase (Table 2). The reason for this seems to be the need to hydrate the Nε of the doubly protonated His which points toward the Trp. The water that moves to hydrate the Nε pushes the Trp side chain inward, partially opening the gate (Fig. 3). This provides a mechanism for opening the gate only from the outside. This change in the Trp gate is not accompanied by opening of the C-terminus, as revealed by CC distances in Table 2. The opening of the gate at Q2 is not exploited by the outward trajectories due to trapping of the H^+^ at the channel periphery.

**Table 2.** Average distances between Trp indole nitrogens and between C-terminal carbon in adjacent helices in the crystal structure (6BKK) and over 200-ps MD simulations.

|  | NN or <NN> | CC or <CC> |
| --- | --- | --- |
| 6bkk 4mer 1 | 4.8 | 17.7 |
| 6bkk 4mer 2 | 4.6 | 17.9 |
| WT Q0 | 5.1 | 17.7 |
| WT Q2 | 6.0 | 17.8 |
| W41F Q0 | - | 17.7 |
| D44N Q0 | 5.1 | 18.0 |
| D44A Q0 | 5.0 | 17.6 |
| W41FD44Q Q0 | - | 17.7 |

**Fig. 3.**
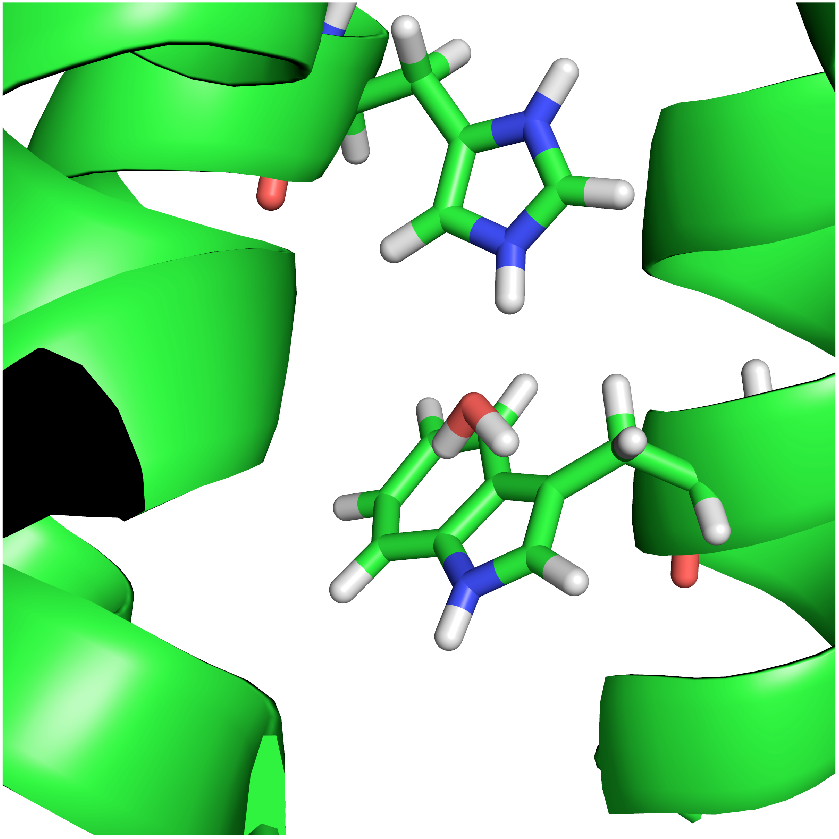
Hydration of the Nε of His pushes the Trp side chain inward.

### Outward crossing probabilities in mutants

Certain mutations of Trp41 and Asp44 are known to allow outward conductance in an outward pH gradient (Tang et al. 2002; Ma et al. 2013). We constructed such mutants by replacing the relevant side chains. 45-ns simulations of the mutants showed them to be stable, with backbone RMS deviations fluctuating below 2 Å (data not shown), except for D44N which deviates beyond 2 Å at ∼40 ns (in previous simulations at Q2 this happened earlier (Ma et al. 2013)). The Trp gate of the Asp44 mutants does exhibit a significant opening compared to the crystal structure. To avoid the uncertainty over the ability of the force field and/or the computational protocol to accurately reproduce such conformational adaptations, we chose to perform hopping simulations near the crystal structure. In short-time-scale simulations there is no significant change in the opening of the channel or the packing of the Trp gate where it exists (Table 2).

Table 3 shows the outward crossing probability for the wild type and mutants at gradually increasing V. Looking at the Q0 trajectories and Table 3 we can identify three barriers for outward H^+^ movement. The highest barrier is escape from Asp44 and starts to be overcome around 8 V. The second barrier is the Trp gate and is overcome around 6.5 V. The third barrier is lack of hydration below the His layer, and is mostly overcome at about 5 V, although it can also stochastically occur even at higher V. Because the Trp gate barrier is lower than the Asp barrier, we do not observe much difference between wild type and W41F.

**Table 3.** Probability of outward H^+^ movement in M2 wild type and mutants at His charge 0 or 2 and different voltages in 100-ps simulations.

|  | Q0-4V | Q0-5V | Q0-6.5V | Q0-8V | Q2-8V |
| --- | --- | --- | --- | --- | --- |
| WT | - | 0% | 0% | 30% | 0% |
| W41F | - | 0% | 20% | 20% | 0% |
| D44N | - | 0% | 60% | 80% | 0% |
| D44A | - | 0% | 40% | 80% | 0% |
| W41F/D44A | 0% | 60% | 60% | 80% | 0% |

In the unsuccessful W41F trajectories the H_3_O^+^ stays bound to Asp or escapes from one to bind to another. Its orientation is mostly H-outward, but sometimes it is also vertical or H-inward. One or two hops separate it from water between the Phe side chains, but these hops are apparently too costly energetically. In addition, there is a dry area above the Phe layer.

The barrier function of the Trp gate is most clearly seen in the Q0-5V results for D44N/A where, in the absence of the Asp barrier, the Trp gate reduces the crossing probability from 60% to 0%. Looking at the D44A/N trajectories we see that there is typically one water molecule between the Trp (“interstitial”) and the H^+^ stays one or two hops away from this water. The immediate neighbor of H_3_O^+^ seems to accept a weak H bond from a Trp NH and that disfavors its protonation (Fig. 4). In other snapshots, the H_3_O^+^ is oriented H-inward and is not even H-bonded to the interstitial water. When Trp is replaced by Phe in the double mutant there is more space and more water between the Phe side chains so that crossing is less unfavorable.

**Fig. 4.**
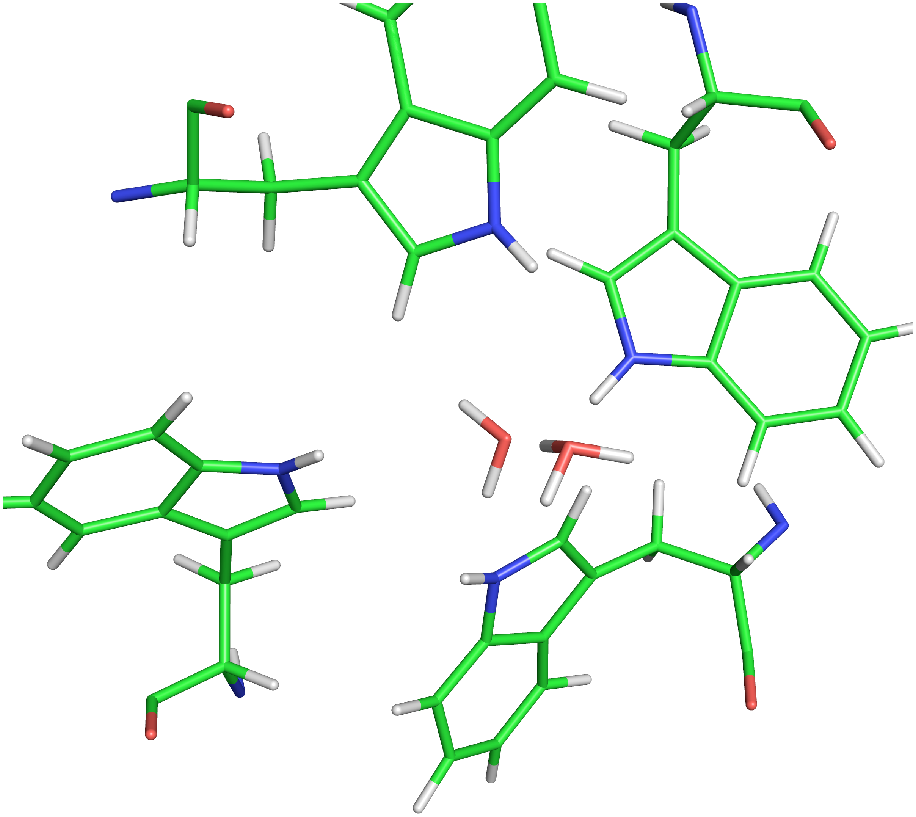
Donation of a proton to the interstitial water between the Trp is energetically costly because that water accepts a weak H bond from Trp.

In the Q2 state the outward crossing probability of WT and all mutants is 0. Undoubtedly, this is caused by His repulsion. However, the same repulsion also exists in the inward direction but does not prevent crossing. This points to an additional feature of the M2 channel that creates asymmetric conductance. Examination of the outward Q2 trajectories of the double mutant revealed that, although there is wide open space between the Phe, the proton tended to reside at the periphery of the inner vestibule (movie S4). The reason for this may be a combination of His repulsion and the known affinity of the excess proton for the water surface and hydrophobic interfaces (Petersen et al. 2004; Iuchi et al. 2009; Lee and Tuckerman 2009; Duignan, Parsons, and Ninham 2015). Notably, the outer and inner vestibules in M2 are very different. The outer vestibule is lined by protein (Ser, Gly, Ala). Because of the conical shape, the inner vestibule is more spacious and has fenestrations covered by lipids. H_3_O^+^ tends to be attracted to these fenestrations, below the Trp or Phe, especially when the His ring is charged. We might call this a hydrophobic trap, in contrast to the electrostatic trap of Asp44. On the channel periphery, as one moves outward there are fewer water molecules, which makes it more difficult to stabilize the positive charge. This may explain the inward rectification (lower current outward than inward at opposite voltages) observed for the D44A mutant, despite the fact that it allows outward pH-driven proton conductance (DiFrancesco et al. 2014).

Other possible origins for the asymmetry in the W41F/D44A mutant were examined and discounted. The first was the tautomer and conformation of the deprotonated His side chains. The HSE (or τ) tautomer chosen initially, points the NH inward. However, replacing HSE by HSD made no difference in the crossing probability. Another possibility was that the repulsion by the Arg45 residues pushed H_3_O^+^ off-center. However, mutation of Arg45 to Ala did not improve conductance but actually reduced it to 0%. The helix dipole could also contribute to directional conductance (López et al. 2012) but this notion is difficult to test (the peptide termini are already neutral in the present simulations).

### H_3_O^+^ orientation around the Trp barrier

Extensive free energy calculations showed that the largest barrier to H^+^ translocation in the M2 channel is at the Trp layer (Liang et al. 2014, 2016). In the outward hopping trajectories we often observed the H_3_O^+^ to be oriented with the H inward. The orientation of H_3_O^+^could affect the translocation ability of the excess proton, and voltage could affect its orientation. Therefore, we performed standard simulations without voltage and proton hops, restraining the hydronium to be between the His37 and Trp41 layers (z=8 Å), or between the Asp44 and the Trp41 layers (z=12 Å), and calculated µ_z_, the component of the dipole vector parallel to the channel axis. Between His and Trp <µ_z_> was -1.1 ± 0.6 for Q0, 1.4 ± 0.3 for Q1, and 1.4 ± 0.6 for Q2. That is, when the His are neutral, H_3_O^+^ is oriented with the H outward, but when the His are charged its orientation changes to H inward, as expected. Between Trp and Asp <µ_z_> was 1.2 ± 0.4 for Q0, 1.2 ± 0.6 for Q1, and 0.7 ± 0.5 for Q2, that is, H_3_O^+^ is oriented H-inward (Fig. 5). At Q2 the repulsion from the His pushes the H_3_O^+^ to the periphery of the channel, where it can orient more freely, and that reduces orientational polarization. The primary reason for the H-inward orientation is interaction with the Asp layer and, when present, the His layer charge. The Trp side chain indole NH groups could also make a contribution. When voltage is applied on the H_3_O^+^ and the surrounding water molecules and hops are allowed, a variety of orientations are observed and this is likely related to the ability of H^+^ to translocate outward when driven by voltage.

**Fig. 5.**
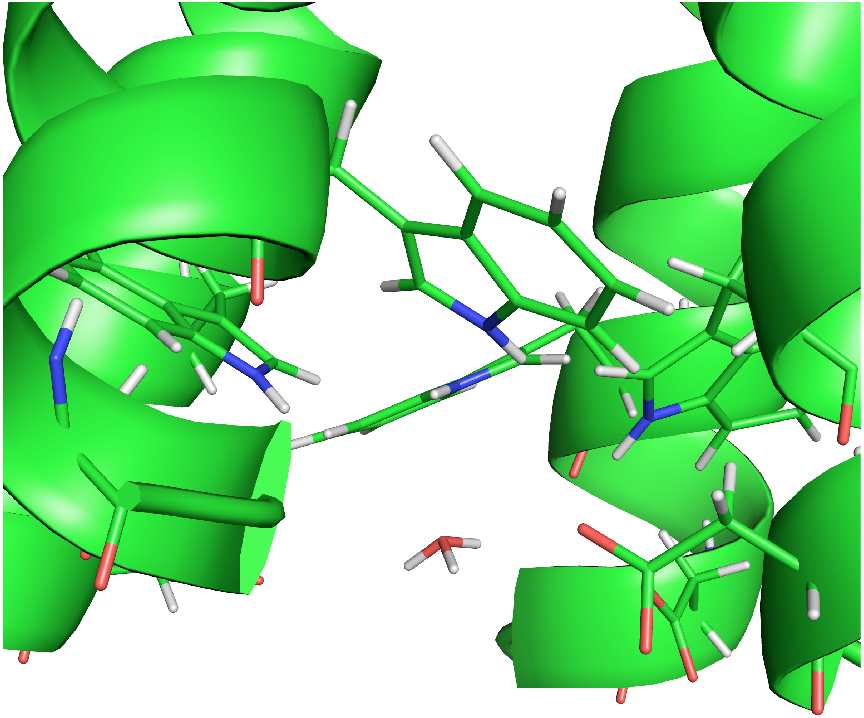
In the absence of voltage, H_3_O^+^ is oriented H-inward between the Asp and Trp layer. Snapshot from Q0 simulation restraining the H_3_O^+^ at z=12Å.

## DISCUSSION

A number of hypotheses have been proposed to explain the inability of M2 to conduct protons outward upon acidification of the interior. The first was that the Trp side chains provide a steric block during outward conduction but a conformational change opens them during inward conduction (Tang et al. 2002). The importance of Asp44 was attributed to its ability to maintain the Trp gate in a closed conformation (Ma et al. 2013). Simulations of the D44N mutant observed a more open and hydrated structure, that was proposed to facilitate outward conductance (Watkins, DeGrado, and Voth 2022). The relative magnitude of inward and outward barriers were also used to provide a kinetic explanation of asymmetric activation and conductance (Liang et al. 2016). The present simulations add three new mechanisms: a) electrostatic trapping by Asp44, b) orientation of H_3_O^+^ with the protons inward, favoring inward conduction, and c) a conical shape which, when combined with repulsion from the His ring, pushes and traps the H_3_O^+^ to the hydrophobic lining of the inner vestibule. It would be worth exploring whether these mechanisms are exploited in other systems that exhibit vectorial proton transport, such as proton pumps.

Orientation of H_3_O^+^ is key in this problem. One-dimensional PMFs of an excess proton along the M2 channel provide valuable information (Liang et al. 2014, 2016; Watkins, DeGrado, and Voth 2022) but neglect the directional nature of Grotthus hopping. Standard transition state theory assumes equal probability for a molecule at the top of the barrier to go forward or backward. However, this is not true for Grotthus hopping. Under the situation depicted in Fig. 6, a H_3_O^+^ would have a much increased probability of crossing the same hydrophobic barrier from the left than from the right. As a consequence, the effective barrier from the left is significantly lower than that from the right. In addition, because of proton hopping, the H_3_O^+^ from the left does not even have to reach the top of the barrier, it just needs to be close enough to donate its proton to a water on the other side.

**Fig. 6.**
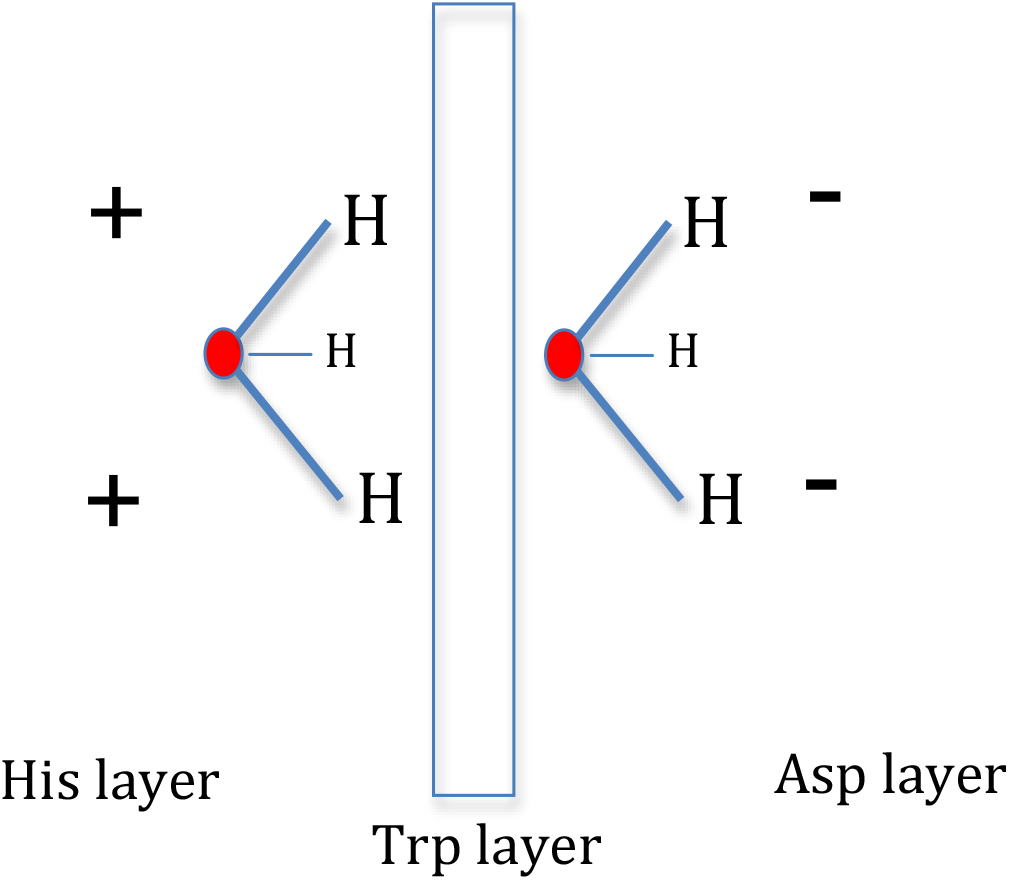
Hydronium is oriented by the surrounding charges interacting with its own partial charge distribution (oxygen negative, protons positive). Its orientation affects the probability of the proton to move left or right from the top of a barrier because Grotthus hopping occurs in the direction to which the H^+^ point.

Differences in conformation could contribute significantly to differences in activity between the mutants. Indeed, NMR experiments in detergent micelles indicated changes in the Trp gate in the D44N mutant and an opening of the structure has been observed in previous simulations (Ma et al. 2013; Watkins, DeGrado, and Voth 2022). However, without full experimental structures for the mutants we would have to rely on the force field and long simulations to produce a structural model. In the present work we chose to leave this factor out of consideration and study proton transport near the experimental wild type structure. The contribution of conformational adaptations could be explored in the future by hopping simulations on MD-relaxed structures and/or experimentally determined mutant structures if they become available.

It is experimentally established that, in contrast to proton concentration, voltage is able to drive outward conductance in the wild type M2 channel (Tang et al. 2002). Based on this, Tang et al. proposed the term gating, rather than rectification, although some genuine inward rectification in the IV curves is also observed (Chizhmakov et al. 2003; DiFrancesco et al. 2014). How can voltage drive outward proton current while proton gradient cannot? These two driving forces differ in a number of ways. Voltage is a long range force, acting directly on H_3_O^+^ and every other charge. It could cause subtle conformational adaptations and affect the protonation and deprotonation rate of His residues. One action of positive voltage we have observed here is to orient the H_3_O^+^ and the waters surrounding it with the protons outward, facilitating outward conduction. The energy change upon flipping a single H_3_O^+^ or H_2_O molecule is very small at values like 60 mV typically used in experiments, but when applied on a large number of molecules over long time scales it could have a significant effect.

In this work the Asp44 appear to be a stronger barrier than the Trp gate by acting as electrostatic traps, whereas experimentally the Trp is known to be at least as important. This could be due to force field inaccuracies, but other explanations are also possible. One is that transient protonation of the Asp44 at low pH could reduce or eliminate their trapping function. The present work was done with only one excess proton. If more than one protons are available, one or two could transiently protonate the Asp44 and the others could then proceed more easily to the Trp gate. Preliminary simulations with four excess protons allowing Asp protonation led to protonation of all four Asp44. This will be investigated more extensively in future studies. More experimental work on the Asp44, such as pKa measurements or proton exchange rates by NMR (Hu, Schmidt-Rohr, and Hong 2012), would also be valuable. Finally, the one-dimensional voltage bias we are using may affect somewhat differently the different barriers, i.e. it may be more effective for the Trp barrier than the Asp barrier.

We note again that the crossing probabilities we calculate here are not physically measurable quantities, like conductances. They depend on the magnitude of the bias force (voltage) and the length of the simulation. For example, the calculation of 0% crossing probability in the Q2 state in no way means that the conductance will be zero. It is also conceivable that outward conductance could occur by deprotonation of the His, lowering of the His ring charge, and reprotonation from inside. To determine the mechanism one would need to calculate actual time scales of deprotonation vs. time scales of conduction at different Q states.

## METHODS

The starting point in this work was the crystal structure of M2 with amantadine (PDB code 6BKK), which belongs to the “C_closed_” class (Thomaston et al. 2018). The inhibitor was deleted. As in many previous simulation studies (Acharya et al. 2010; Ma et al. 2013; Wei and Pohorille 2013; Dong et al. 2014; Liang et al. 2014; Watkins, DeGrado, and Voth 2022), the M2 tetramer was embedded into a POPC membrane. The membrane had been generated using CHARMM-GUI (Wu et al. 2014) and equilibrated with a different M2 crystal structure which was replaced by 6BKK and reequilibrated for 10 ns keeping the protein fixed. 37 water molecules in the pore lumen from the crystal structure were preserved, replacing an equal number of overlapping waters from the previously equilibrated system. The system contained about 32,000 atoms - the 4 peptide monomers, 138 POPC lipids, and about 4,060 TIP3P water molecules in a 70 Å X 70 Å X 63 Å box. The His 37 were either all singly protonated (tautomer τ, or HSE, with proton on Nε) or had a total charge of +1 or +2. In the latter case two diagonally positioned His residues were doubly protonated. The peptide termini were neutralized, which is more realistic for the full-length peptide, for truncated peptides with blocked termini, and even for unblocked truncated peptides with counterions (no salt is present in the current simulations). Preliminary results (not shown) with charged termini found an even stronger preference for inward proton conductance due to the electric field the termini create. We used the charmm36 force field for the protein (Best et al. 2012), a timestep of 2 fs, a cutoff of 12 Å for the van der Waals interactions, and Particle Mesh Ewald (Essmann et al. 1995) for long range electrostatics. The simulations were run at a constant temperature of 303 K and a constant pressure of 1 bar using the Nose-Hoover and Langevin piston methods. Mutant structures were generated by replacing the Asp44 and/or Trp41 side chain, building the unknown coordinates of the new side chain in a standard geometry, and energy-minimizing.

The excess proton was modeled as a classical hydronium (Sagnella and Voth 1996). The proton hopping simulations were carried out with the MOBHY module (Lazaridis and Hummer 2017) implemented in a custom version of the program CHARMM (Brooks et al. 2009). Grotthus hopping is enabled by periodic attempts (here every 10 steps) to move a proton from the current H_3_O^+^ to a water molecule that is hydrogen bonded to it. The hop is accepted when the energy change is less than a threshold, here 20 kcal/mol, which was empirically chosen to reproduce the proton diffusivity in water. In case of hop acceptance, the geometry of the two partners is optimized for 40 steps, which approximately removes the energy that the hop introduces into the system. In this method, a number of TIP3P residues (here 15) are replaced by H3O residues with a dummy H (effectively identical to TIP3P) and hops occur between the actual H_3_O^+^ and these H3O residues. When proton donation to TIP3P is considered, the TIP3P is switched with an H3O residue. Harmonic restraints were used to keep the membrane-protein system at the center of the box, the protein at a certain point in the xy plane, and the H3O residues in a 5-Å cylinder around the channel axis.

Because proton conduction through the M2 channel is a slow process (Mould et al. 2000; Hong and DeGrado 2012), it needs to be accelerated in some artificial way. For that we used voltage, which was applied in a 55-Å wide region across the membrane (Mottamal and Lazaridis 2006) and felt only by the hydronium and the surrounding H3O residues (otherwise the membrane would be destabilized by the high voltage used). The magnitude of the voltage was chosen empirically to be just large enough to start seeing differences in conductance inward and outward or between different mutants. The same bias is applied inward and outward, so it is expected to preserve the relative facility of movement in the two directions. However, it is not possible to recover the correct kinetics and the true absolute conductance from these simulations. Although we will use the word “conductance” as equivalent to crossing probability, it should be understood that we are not making rigorous conductance estimates, but only compare inward and outward mobility at the same biasing force. All proton hopping simulations lasted 100 ps. Although it would be possible to use smaller bias and longer simulations, the distribution of first passage times at smaller biases would be broader and it would be more difficult to observe statistically significant differences. When His titration is allowed, the parameters for “fast” His (Lazaridis and Hummer 2017) were used to artificially accelerate protonation and, especially, deprotonation.

## Acknowledgments

Support by the National Science Foundation (MCB-1855942) is gratefully acknowledged.

## Supporting Information

A sampling of movies of proton hopping trajectories. The full trajectories corresponding to these movies will be deposited at zenodo.org.

